# Computing personalized brain functional networks from fMRI using self-supervised deep learning

**DOI:** 10.1101/2021.09.25.461829

**Authors:** Hongming Li, Srinivasan Dhivya, Zaixu Cui, Chuanjun Zhuo, Raquel E. Gur, Ruben C. Gur, Desmond J. Oathes, Christos Davatzikos, Theodore D. Satterthwaite, Yong Fan

**Affiliations:** Center for Biomedical Image Computation and Analytics, University of Pennsylvania, Philadelphia, PA 19104, USA; Department of Radiology, University of Pennsylvania, Philadelphia, PA 19104, USA; Chinese Institute for Brain Research, Beijing, 102206, China; Key Laboratory of Brain Circuit Real Time Tracing (BCRTT-Lab), Tianjin University Affiliated Tianjin Fourth Center Hospital; Department of Psychiatry, Tianjin Medical University, Tianjin, China; Center for Neuromodulation in Depression and Stress, University of Pennsylvania, Philadelphia, PA 19104, USA; Department of Psychiatry, University of Pennsylvania, Philadelphia, PA 19104, USA; Penn Lifespan Informatics and Neuroimaging Center, University of Pennsylvania, Philadelphia, PA 19104, USA; Penn/CHOP Lifespan Brain Institute, University of Pennsylvania, Philadelphia, PA 19104, USA; Department of Neurology, University of Pennsylvania, Philadelphia, PA 19104, USA

## Abstract

A novel self-supervised deep learning (DL) method is developed for computing bias-free, personalized brain functional networks (FNs) that provide unique opportunities to better understand brain function, behavior, and disease. Specifically, convolutional neural networks with an encoder-decoder architecture are employed to compute personalized FNs from resting-state fMRI data without utilizing any external supervision by optimizing functional homogeneity of personalized FNs in a self-supervised setting. We demonstrate that a DL model trained on fMRI scans from the Human Connectome Project can identify canonical FNs and generalizes well across four different datasets. We further demonstrate that the identified personalized FNs are informative for predicting individual differences in behavior, brain development, and schizophrenia status. Taken together, self-supervised DL allows for rapid, generalizable computation of personalized FNs.

## INTRODUCTION

Functional magnetic resonance imaging (fMRI) is a powerful tool to study functional networks (FNs) in the human brain. Accumulating evidence demonstrates that FNs undergo predictable normative development in youth and abnormal development is associated with diverse psychopathology^1^. However, most of our knowledge of FNs is drawn from studies that used “one size fits all” group atlases that are not tailored to the individual. Recent convergent evidence from multiple independent efforts has established that FNs are indeed person-specific. These studies emphasize that there is marked inter-individual variability in the *functional topography* of FNs, which is defined as the location and spatial arrangement of FNs on the anatomic cortex^2–11^. Notably, it has been shown that the greatest variability in functional topography is present in association networks such as the fronto-parietal control network (FPN), the default mode network (DMN), and the ventral attention network (VAN). Notably, these association networks are of particular interest as they undergo remarkable change over the lifespan and are most impacted by psychopathology^5, 11^. Group-level definitions of FNs used in the vast majority of translational studies fail to account for such inter-individual variability of FNs, inevitably mixing signals from disparate FNs, and reducing sensitivity in clinical applications.

An alternative to using group-level atlases is to create personalized FNs^5, 10, 12–22^. To enforce correspondence of FNs across different individuals, these methods typically compute personalized FNs under certain constraints to be similar to group level FNs, such as those built upon independent component analysis (ICA)^20, 21^ or based on assumptions that loadings of corresponding FNs of different individuals follow certain distributions^5, 16, 19, 22^. However, such techniques may yield biased results since little is known about FNs’ statistical distribution. Such limitations have been partially overcome using spatially-regularized non-negative matrix factorization^10^. Recent studies have also demonstrated that deep learning methods, such as deep autoencoders, can be used to learn low-dimensional representations of FNs in a latent space from fMRI data^23–27^. However, these methods have thus far not been used to create personalized FNs.

Since personalized FNs account for inter-individual differences in functional neuroanatomy^28^, we developed a novel self-supervised deep learning (DL) method for computing personalized FNs from fMRI data while maintaining inter-individual correspondence. We hypothesized that learning an effective intrinsic representation of FNs in a low-dimensional latent space would facilitate the computation of bias-free personalized FNs with improved functional homogeneity. We adopted convolutional neural networks (CNNs) with an encoder-decoder architecture to identify individual-specific FNs in an end-to-end learning fashion. In this framework, the encoder is used to learn an effective intrinsic representation of FNs in a low-dimensional latent space and the decoder is used to infer personalized FNs with inter-individual correspondence from the effective representation. We trained the encoder-decoder CNNs using self-supervised information such as functional homogeneity and spatial sparsity; this trained model could then be applied to fMRI data of an unseen individual’s data to identify FNs in one forward-pass computation. We have trained a DL model on resting-state fMRI (rsfMRI) data of individuals from the Human Connectome Project (HCP)^29^, and applied it to rsfMRI data of new individuals from different datasets including healthy individuals with different age ranges and patients with schizophrenia. Results reveal that the DL model could obtain personalized FNs which coincide with well-established large-scale FNs. Furthermore, these personalized FNs were informative for predicting behavioral measures and brain age of the healthy individuals, and also could distinguish schizophrenia patients from healthy individuals. As described below, these results indicate that the personalized FNs identified by our DL method accurately capture the variability of personalized functional neuroanatomy across diverse datasets in a way that meaningfully captured individual differences in behavior and brain health. This proposed method marks a paradigm shift in identifying personalized FNs from matrix decomposition to learning based predictive modeling.

## RESULTS

Our self-supervised DL framework includes a representation learning module and a functional network learning module for identifying personalized FNs (**Fig. 1a)**. The representation learning module (**Fig. 1b**) learns time-invariant feature maps from the input fMRI, and the functional network learning module (**Fig. 1c**) with an Encoder-Decoder architecture computes personalized FNs from these feature maps by optimizing self-supervision measures including functional homogeneity and spatial sparsity so that the network is trained and optimized based on the input fMRI data without any external supervision.

**Fig. 1.**
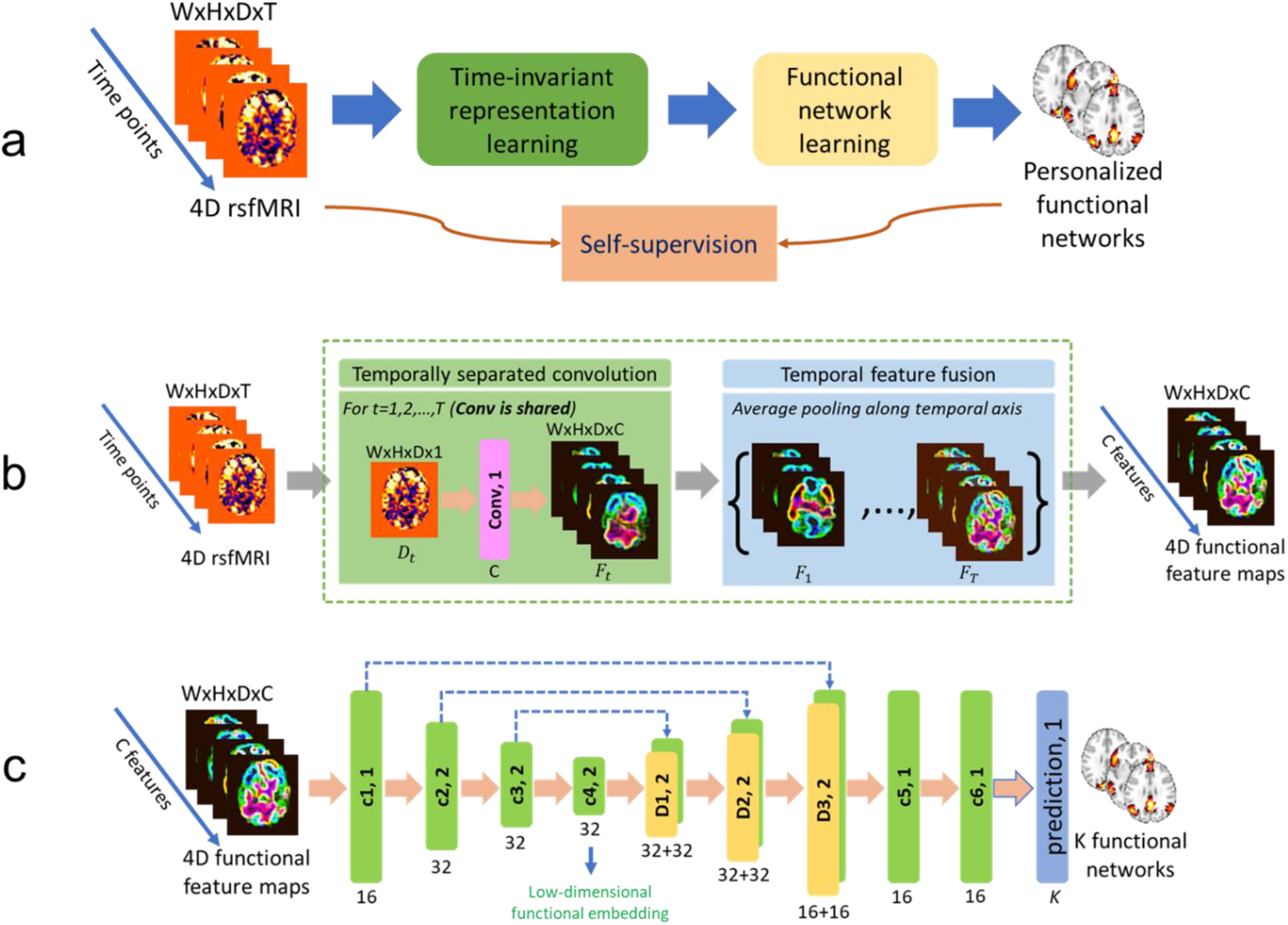
Our proposed deep learning framework for personalized functional networks. **(a)** Schematic diagram of the self-supervised deep learning model for identifying personalized functional networks (FNs). **(b)** Network architecture of the time-invariant representation learning module to account for temporal misalignment of resting-state fMRI data across different scans. **(c)** Network architecture of the functional network learning module for the prediction of personalized FNs. The numbers underneath convolutional (Conv, c1, c2, c3, c4, c5, and c6) and deconvolutional (d1, d2, and d3) layers indicate their corresponding numbers of kernels with a stride of 1 or 2. The kernel size in all layers is set to 3×3×3.

### Self-supervised DL model could identify FNs which coincide with well-established FNs

We trained a DL model (**Fig. 1**) on rsfMRI data of 400 individuals randomly selected from the HCP cohort. The number of FNs was set to 17 in order to facilitate a direct comparison with well-established FNs^30^. The trained DL model was applied to another 678 HCP individuals served as testing data for evaluation. As two 3T rsfMRI sessions were acquired for each individual, we have adopted the first session as the primary dataset (HCP REST1) and used the second session for replication (HCP REST2). The personalized FNs of all testing individuals computed by the DL model passed the sanity testing with respect to both functional homogeneity and spatial correspondence. Particularly, the personalized FNs had higher within-network functional homogeneity than the group average FNs of all testing individuals and maintained spatial correspondence with their corresponding group average FNs (**Fig. S1**). The average FNs of the 678 testing individuals identified by our DL model on the REST1 dataset are illustrated in **Fig. 2**. It can be observed that the DL model successfully identified both spatially localized and distributed FNs, including visual networks, somatomotor networks, dorsal attention networks (DAN), ventral attention networks (VAN), fronto-parietal networks (FPN), default mode networks (DMN). These identified FNs show high correspondence to the well-established functional atlases^30^. Using a conservative spatial permutation test based on spatial-correlation-preserving surrogate maps generated by BrainSMASH^31^, we found a significant alignment (*p* < 0.001) between the FNs identified by the DL model and the canonical networks from the Yeo atlas^30^.

**Fig. 2.**
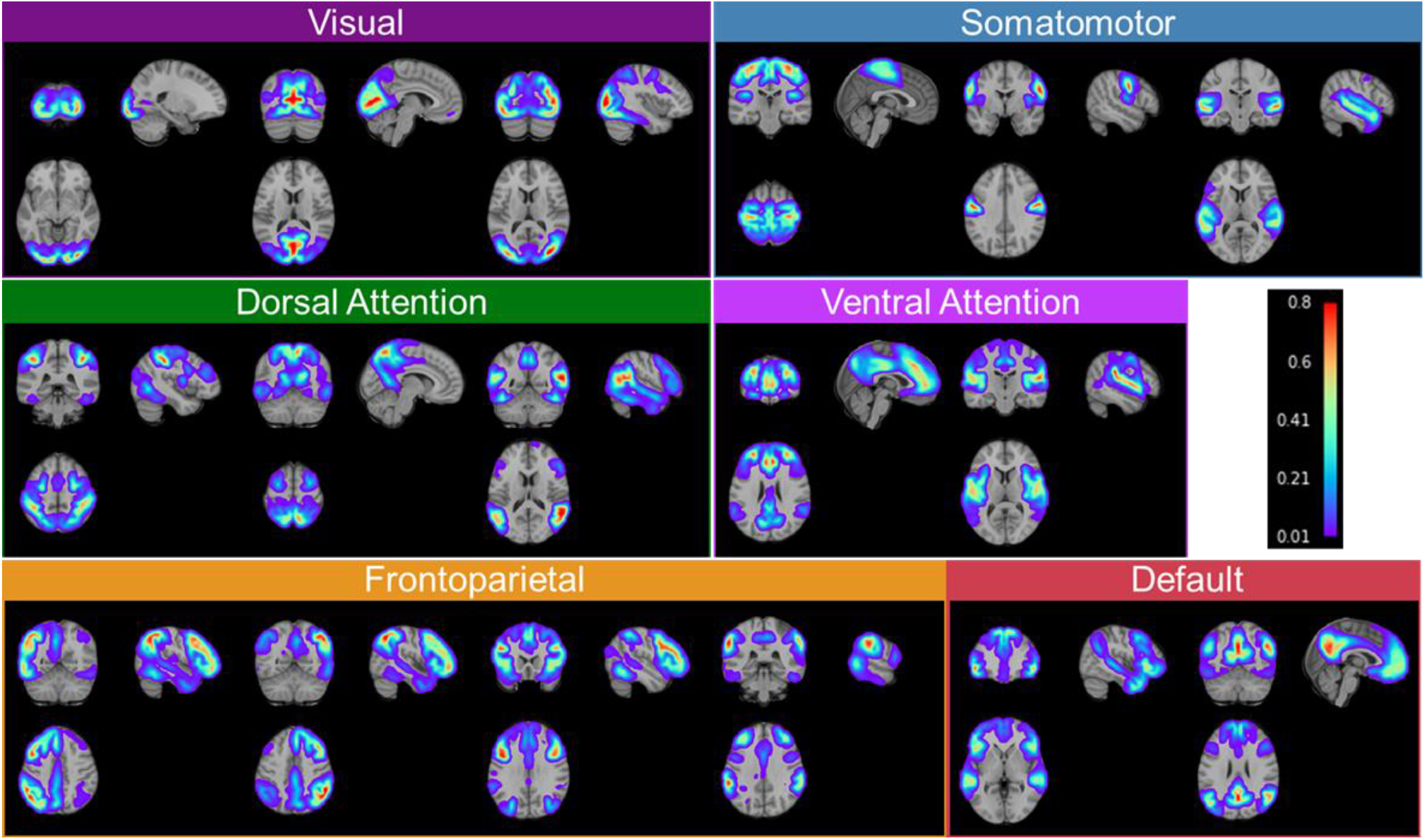
Group average functional networks identified by the DL model on the testing individuals of HCP REST1 dataset. The 17 networks have been categorized into visual networks, somatomotor networks, dorsal attention networks, ventral attention networks, fronto-parietal networks, and default mode networks, based on their spatial overlap with the Yeo atlas^30^.

### Self-supervised DL model could identify personalized FNs with improved functional homogeneity

Personalized FNs of three randomly selected testing individuals from HCP REST1 dataset are illustrated in **Fig. 3a**. Inter-individual differences can be clearly observed in the personalized FNs, illustrating that the proposed method captured the inherent differences of individualized functional neuroanatomy. We also compared the personalized FNs computed using the proposed DL model and a top-performing spatially-regularized NMF method^10^ in terms of their within-network functional homogeneity^10^. The personalized FNs obtained by the DL model had significantly higher functional homogeneity than those computed using the spatially-regularized NMF (**Fig. 3b**, *p* < 10^−5^, Wilcoxon signed rank test). This increase in functional homogeneity may happen because no specific regularization was adopted to enforce the FNs to follow a prior distribution – facilitating better characterization of inter-individual variability in functional neuroanatomy.

**Fig. 3.**
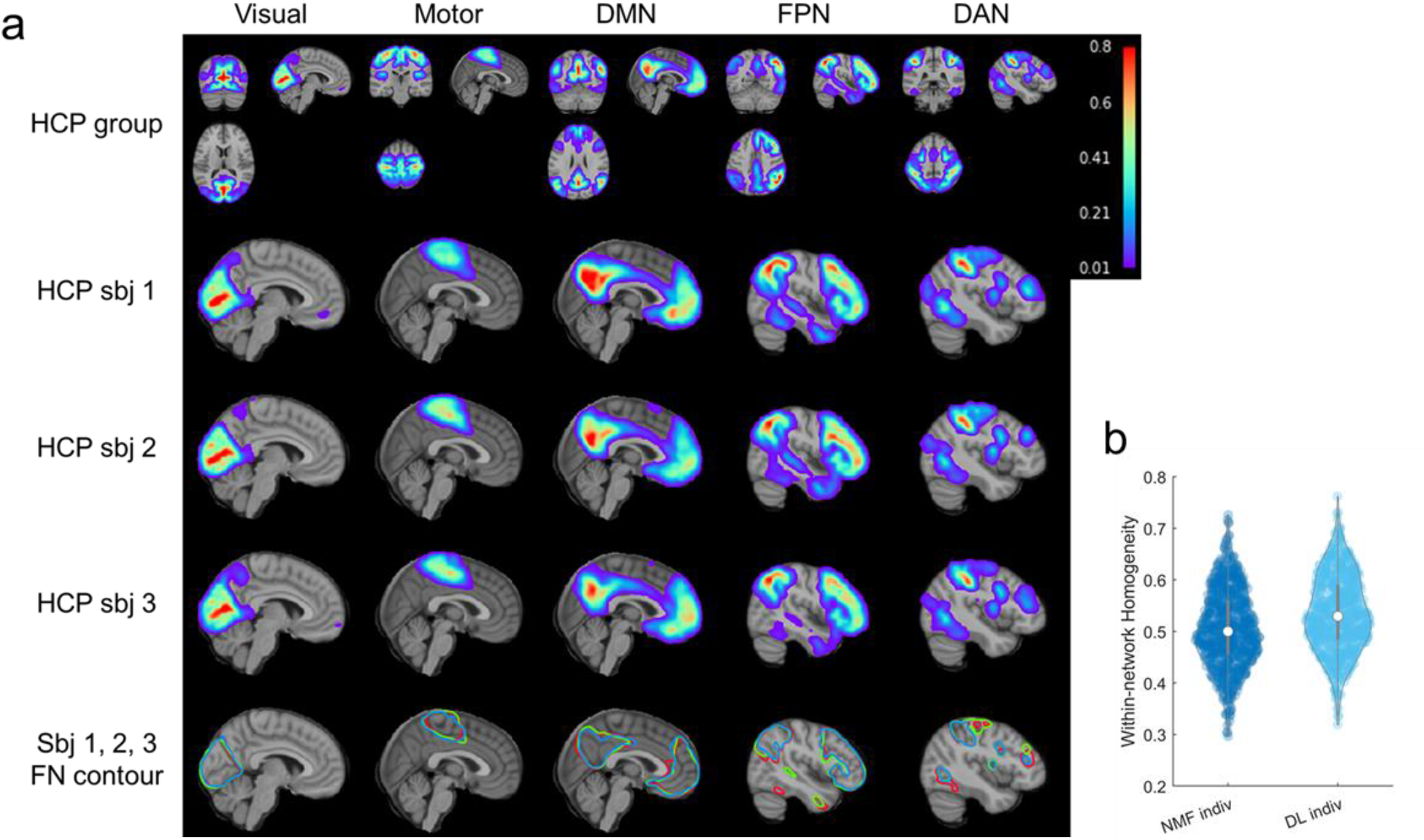
Personalized functional networks identified by the DL model for three randomly selected individuals from the testing cohort of HCP REST1 dataset, including visual network, somatomotor network, default mode network (DMN), fronto-parietal control network (FPN), and dorsal attention network (DAN). (**a**) Average FNs of all testing HCP subjects (the first row), three individual subjects’ personalized FNs in sagittal view (the second, third, and fourth rows), and isolines of the FNs at a value of 0.15 in different colors (the fifth row). (**b**) Personalized FNs computed using the DL model had significantly higher functional within-network homogeneity than those computed using the spatially-regularized NMF (*p* < 10^−5^, Wilcoxon signed rank test). Each datapoint in the plot denotes the average functional homogeneity across all FNs of one subject.

### Personalized FNs from the DL model are highly reproducible

To evaluate the reproducibility of personalized FNs computed by the DL model, we compared personalized FNs of the same testing subjects, computed from their rsfMRI scans collected in different imaging sessions, including the HCP REST1 (R1) and REST2 (R2) datasets. Default mode network and fronto-parietal network of three randomly selected testing individuals are shown in **Fig. 4a**, illustrating that the intra-subject (inter-session) differences between corresponding FNs are visually smaller than the inter-subject counterpart. To evaluate this quantitatively, we further tested if the personalized FNs computed from two imaging sessions (R1 and R2) of the same subjects are more similar than those of different subjects so that FNs can be used to identify subjects. As illustrated in **Fig. 4b**, each subject in R1 was compared with all subjects in R2 to identify the individual with the maximally similar FNs. Average spatial correlation coefficient across FNs was used as the similarity metric. The subject identification was considered as correct if the maximally similar FNs between the two sessions were from rsfMRI scans of the same subject. The identification rate was calculated as the ratio between the number of correctly identified target individuals and the total number of target individuals. The identification procedures were carried out in two directions, i.e., identification of subjects of R1 from R2 (R1->R2) and vice versa (R2->R1). The identification rate was 93.6% and 97.4%, respectively for the procedures of R1->R2 and R2->R1 when all 17 FNs were used for the identification (**Fig. 4c**), which was similar to that obtained by the functional connectome fingerprinting study^32^. Since the inter-subject functional topography variability is maximal in association networks^4, 5, 11, 33^, the same subject identification procedure was also carried out based on the fronto-parietal FNs alone (as illustrated in **Fig. 2**), and improved identification rates were obtained, with 98.3% and 98.8% respectively for the procedures of R1->R2 and R2->R1 (**Fig. 4c**). The subject identification performance is on par with that obtained on the HCP subjects in the functional connectome fingerprinting study^32^, although our study used a much larger testing dataset. These results indicate that the DL model could identify reproducible and subject-specific FNs that capture variations of inter-subject functional neuroanatomy.

**Fig. 4.**
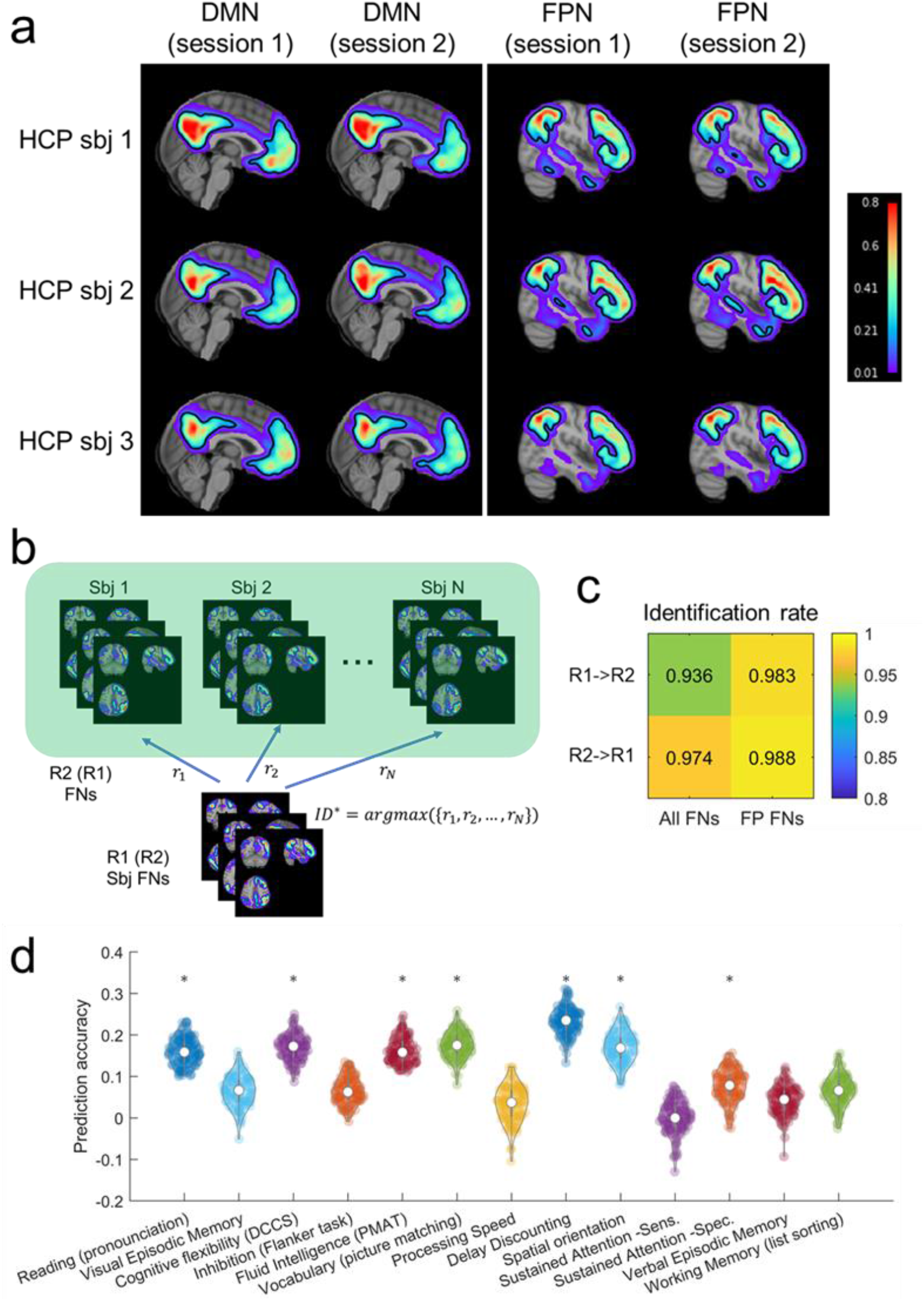
Personalized functional networks (FNs) identified by the DL model are with high reproducibility and associated with behavior. **(a)** Personal FNs including DMN and FPN of three randomly selected individuals using HCP REST1 and REST2 rsfMRI data. The isoline of value 0.15 in each FN is demonstrated by the black contour to facilitate the visual comparison across sessions and subjects. **(b)** FN based identification procedure. Given a query set of FNs from one target subject (sbj), we computed the similarity *r_i_* between its FNs and all the sets of FNs in the Database. The matched identity *ID*\* is the one with the highest similarity. **(c)** The identification rate when 17 FNs (All FNs) or combined fronto-parietal FNs (FP FNs) were used for the identification. **(d)** Prediction accuracy of 13 cognitive measures based on personalized FNs computed by the proposed DL model. Violin plots show the distribution of prediction accuracy of 100 repetitions for each measure. Asterisks denote the cognitive measures that are significantly associated with personalized FNs.

### Personalized FNs from DL model are associated with behavior

To determine whether the individual variations in FNs are behaviorally meaningful, we investigated whether personalized FNs identified by the DL model could be used to predict individuals’ behavior and cognitive performance, with a focus on 13 cognitive measures highlighted in the HCP data dictionary^5^. The loading coefficients of all 17 FNs were concatenated into a feature vector for each individual, and ridge regression with 2-fold cross-validation was used to predict the behavior measures respectively. The regularization parameter in ridge regression was optimized using nested 2-fold cross-validation. Partial correlation coefficient between the predicted and real measures was adopted to evaluate the prediction accuracy, with age, sex, and in-scanner motion as co-variates. The prediction for each behavior measure was repeated 100 times. As shown in **Fig. 4d**, the cognitive measures were predicted with an average accuracy of 0.1092 ± 0.0097 (7 out of 13 were significant, *p* < 0.05, permutation test), in par with the prediction performance obtained in a previous study^5^. These results indicated that the DL model was able to identify the variability of personalized FNs that meaningfully captured individual differences in behavior.

### Self-supervised DL model robustly generalizes to new datasets

To determine whether the DL model was generalizable to individuals whose fMRI scans were collected with imaging protocols/parameters different from the fMRI data used to train the DL model, we applied the model built on the HCP training cohort to fMRI data from two external datasets. Notably, both datasets were different from the HCP cohort in terms of age, scan length, and scanning protocol. The first cohort contained 969 individuals (ages from 8 to 23) from the PNC^34^, and the second cohort included 210 individuals (ages from 3 to 20) from the PING cohort^35^. The personalized FNs of all those individuals computed by the DL model passed the sanity testing with respect to both functional homogeneity and spatial correspondence (**Fig. S1**). Five representative personalized FNs of three randomly selected individuals from the PNC and PING cohort are shown in **Fig. 5a** and **Fig. S2a**, respectively. It can be observed that the DL model generalized well to individuals from different cohorts, with both localized and distributed FNs identified successfully. Differences in FNs could also be clearly observed across individuals for both cohorts. The FNs obtained by the DL model had significantly higher functional homogeneity than those computed using the spatially-regularized NMF (**Fig. 5b** and **Fig. S2b**, *p* < 10^−5^, Wilcoxon signed rank test).

**Fig. 5.**
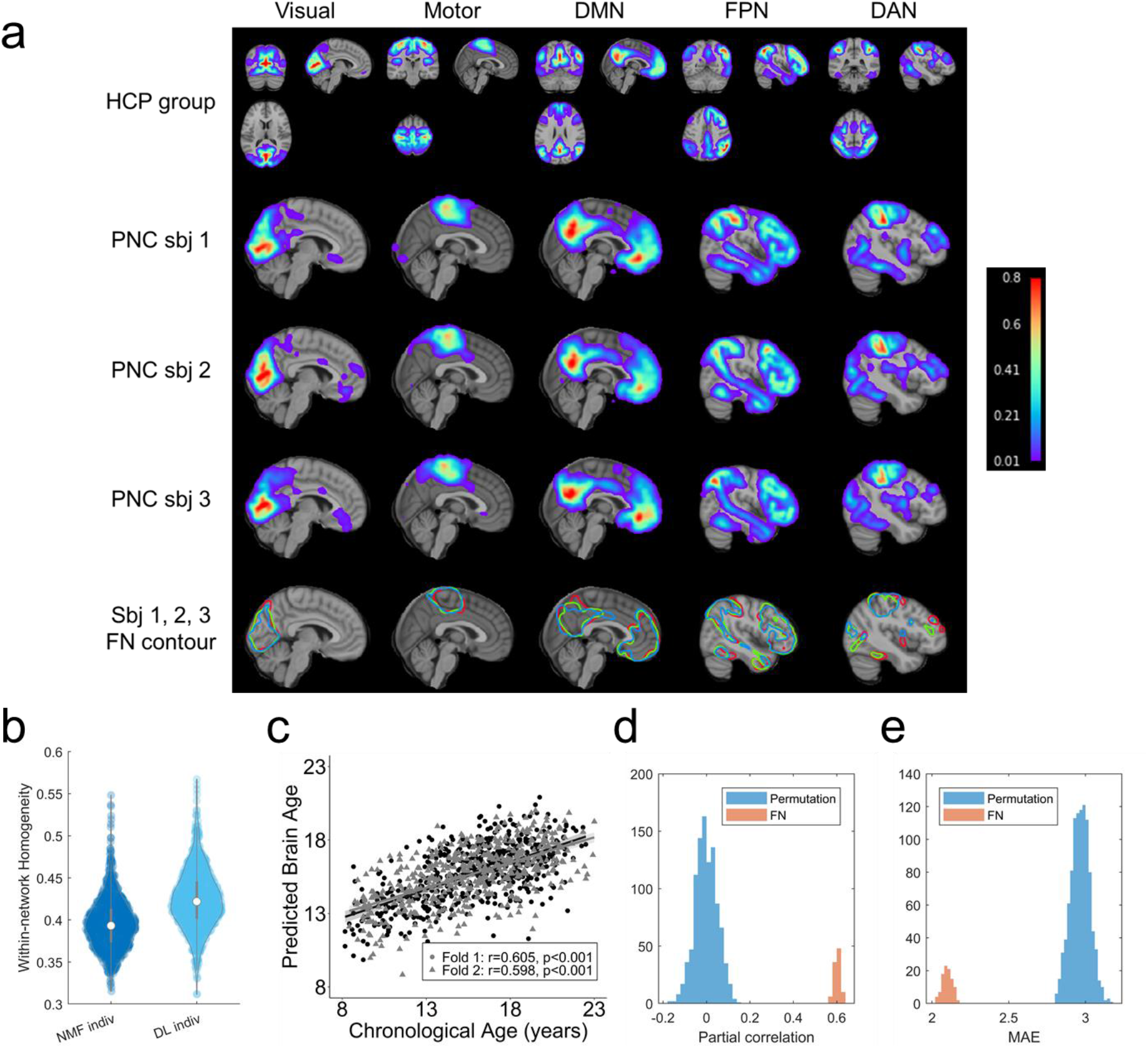
The DL model trained on the HCP cohort generalizes well to the PNC. **(a)** Personalized FNs of three randomly selected PNC subjects, identified by the DL model trained on HCP cohort, in sagittal view (the second, third, and fourth rows), isolines of the FNs at a value of 0.15 in different colors (the fifth row), and their corresponding average FNs of all testing HCP subjects (the first row). **(b)** Personalized FNs computed using the DL model had significantly higher functional within-network homogeneity than those computed using the spatially-regularized NMF (*p* < 10^−5^, Wilcoxon signed rank test). **(c)** Age prediction performance (partial correlation coefficients) of the personalized FNs, obtained with one run of the 2-fold cross-validation. Data points in different colors represent the prediction results of different folds in the 2-fold cross-validation. **(d, e)** Prediction accuracy measured with partial correlation coefficients and MAE of 100 runs of the 2-fold cross-validation and the null distribution of prediction accuracy from permutation test (1000 runs of 2-fold cross-validation using permuted data).

To further determine whether these personalized FNs are biologically meaningful, we investigated whether personalized FNs identified by the DL model could be used to predict individuals’ age, as a recent study demonstrated that functional topography could encode development^11^. The same strategy as the prediction of behavior phenotypes on HCP cohort was adopted for the age prediction. The prediction accuracy was measured with partial correlation coefficient between the predicted brain age and chronological age with sex and in-scanner motion as co-variates, in addition to mean absolute error (MAE). The prediction results shown in **Fig. 5c** and **Fig. S2c** revealed that the personalized FNs identified by the DL model could predict age accurately on both cohorts, with an average partial correlation coefficient of 0.6014 ± 0.0153 (*p* < 0.001, permutation test) and *MAE* = 2.099 ± 0.0324 (*p* < 0.001, permutation test) on the PNC (**Fig. 5d** and **Fig. 5e**), and an average partial correlation coefficient of 0.7838 ± 0.0168 ( *p* < 0.001, permutation test) and *MAE* = 2.9201 ± 0.0861 ( *p* < 0.001, permutation test) on the PING cohort (**Fig. S2d** and **Fig. S2e**). Together, these results indicate that the DL model generalizes well to new datasets.

### Self-supervised DL model could capture the differences of FNs between healthy control and schizophrenia patients

The DL model trained on HCP cohort generalized well to the PNC and PING cohort, despite they contain subjects of quite different age ranges. To determine whether the DL model trained on healthy individuals (HCP cohort) is applicable to individuals with neuropsychiatric illness, we further applied it to one additional dataset, consisting of fMRI data from 101 healthy controls (HC) and 94 schizophrenia (SCZ) patients. The personalized FNs of all those individuals computed by the DL model passed the sanity testing with respect to both functional homogeneity and spatial correspondence (**Fig. S1**). Five representative personalized FNs of three randomly selected individuals (2 HC and 1 SCZ) are shown in **Fig. 6a**, suggesting that the DL model could identify personalized FNs for both healthy controls and patients, with significantly higher functional homogeneity than those computed using the spatially-regularized NMF (**Fig. 6b**, *p* < 10^−5^, Wilcoxon signed rank test).

**Fig. 6.**
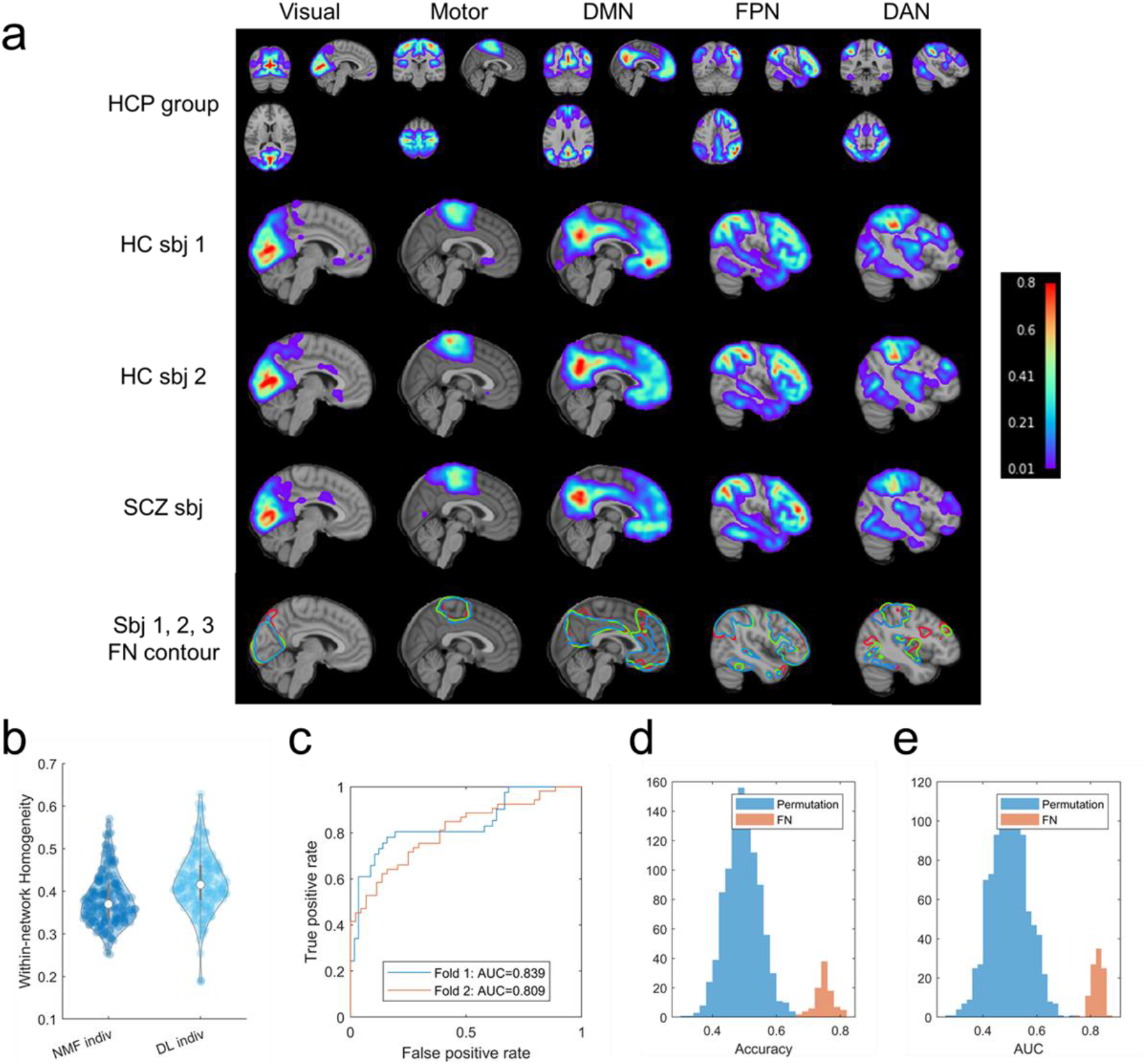
The DL model trained on HCP cohort could capture the differences of FNs between healthy controls (HC) and schizophrenia (SCZ) patients. **(a)** Personalized FNs of two HCs (the second and third rows) and one SCZ patient (the fourth row), identified by the DL model trained on HCP cohort, isolines of the FNs at a value of 0.15 in different colors (the fifth row), and their corresponding average FNs of all testing HCP subjects (the first row). **(b)** Personalized FNs computed using the DL model had significantly higher functional within-network homogeneity than those computed using the spatially-regularized NMF (bottom right, *p* < 10^−5^, Wilcoxon signed rank test). **(c)** Receiver operating characteristic (ROC) curves of classification models built on the personalized FNs of one run of the 2-fold cross-validation. **(d, e)** Classification rates and AUC values of 100 runs of 2-fold cross-validation, and the null distribution of classification rates and AUC values from permutation test (1000 runs of 2-fold cross-validation using permuted data).

To further determine whether the differences of FNs between HC and SCZ patients can identify features relevant to neuropsychiatric illness, we investigated whether the personalized FNs identified by the DL model could be used to distinguish HCs from patients with SCZ in a pattern classification setting. Support vector machine (SVM) with linear kernel was used for the classification with the loading coefficients of all 17 FNs used as features. The classification was carried out under a 2-fold cross-validation setting for training and testing, and this procedure was repeated 100 times. The prediction performance was evaluated with classification accuracy and area under the Receiver operating characteristic (ROC) curve (AUC). The classification based on personalized FNs distinguished HCs from SCZ patients (**Fig. 6c**), with an average classification rate of 0.7503 ± 0.0283 (*p* < 0.001, permutation test) and an average AUC value of 0.8245 ± 0.0204 (*p* < 0.001, permutation test) over 100 classification runs (**Fig. 6d** and **Fig. 6e**). These results reinforce that the DL model generalizes well and is sensitive to individual differences in functional neuroanatomy associated with schizophrenia.

### Self-supervised DL model could identify personalized FNs rapidly

The computational time to compute FNs using our trained DL model was proportional to the scan length of the fMRI scan. On average, it took 22.5 ± 3.1, 4.8 ± 0.2, 7.14 ± 2.61, and 4.26 ± 0.645 seconds to compute 17 FNs for each testing individual subject of the HCP, PNC, PING, and SCZ datasets respectively, highlighting the DL model’s computational efficiency.

## DISCUSSION

A rapidly accruing body of evidence has demonstrated that the spatial distribution of FNs on the anatomic cortex differ substantially across individuals, and that personalized FNs are necessary to account for individual variation in functional neuroanatomy^2–11, 28^. Here we present a novel method for identifying the personalized FNs through self-supervised deep learning. Our self-supervised deep learning method is capable of learning an effective intrinsic representation of FNs in a low-dimensional latent space, which captures the differences in functional topography across individuals and therefore facilitates the computation of personalized FNs by optimizing functional homogeneity of the personalized FNs^10, 36^. We demonstrated that our deep learning method could identify well-established FNs at an individual level with high reproducibility, and that the individual variation of FNs is relevant to individual behavior, brain development, and neuropsychiatric illness such as schizophrenia.

The traditional paradigm of identifying personalized FNs is to model them as latent factors of rsfMRI data in an unsupervised learning framework using matrix decomposition techniques. Most of the existing methods adopt a two-step inference strategy that first estimates group FNs based on fMRI data from a group of individuals and then use the group FNs to infer personalized FNs under different assumptions^5, 13, 20, 23, 37, 38^. As a result, these methods inevitably introduce bias to the group FNs and do not necessarily yield an optimal solution since their personalized FNs are not optimized in the same way as their group level counterparts. Several alternative methods have been developed to identify personalized FNs of different individuals jointly and enforce inter-individual correspondence of FNs using spatial group sparsity regularization^10^ or regularizations based on certain assumptions about statistical distribution of loadings of corresponding FNs of different individuals^16, 19^. Deep belief networks (DBNs) have also been utilized to identify FNs of multiple individuals jointly^39^. However, all these methods cannot directly make inference for new individuals and are computationally expensive. CNNs have also been used to predict individual-specific DMN in a supervised learning setting^40^, with DMNs obtained by conventional brain decomposition methods as ground truth. However, CNNs in the supervised learning setting may be limited by the silver standard ground truth adopted.

Our method is different from the traditional paradigm mainly in two aspects. First, our method is under the paradigm of predictive modeling, which learns the personalized FNs from rsfMRI data in an end-to-end self-supervised deep learning fashion. Our deep learning model is trained based on self-supervised information including functional homogeneity and spatial sparsity of FNs, which do not require external guidance such as FNs computed using conventional techniques. Therefore, any rsfMRI data could be used to train the model. Once the training is completed, the model can be used to identify personalized FNs from a new, unseen individual using one forward-pass computation without extra optimization. This feature makes our method very efficient: it takes about 5 seconds to compute the FNs for an fMRI scan with spatial resolution of 3mm and scan length of 120 time points (TR=2 seconds) on one modern NVIDIA GPU. Such computational efficiency facilitates its use in large-scale studies. Second, the inter-individual correspondence of FNs is enforced implicitly though the encoder-decoder architecture of our deep learning model. An effective intrinsic representation of FNs in a low-dimensional latent space is extracted by the encoder from the original functional signal, from which the decoder can infer personalized FNs with inter-individual correspondence. From a perspective of predictive modeling, voxels with similar functional profiles will be assigned to the same FN by the deep learning model. While different output channels of the deep learning model capture different FNs, one output channel will identify FNs across individuals with functional correspondence. Compared with explicit spatial regularization, this kind of implicit enforcement is more flexible to characterize the inter-individual variation and therefore facilitates the identification of FNs with improved functional homogeneity.

The proposed deep learning model generalizes well from the training dataset to different external datasets without fine-tuning, even though the external datasets used had notably different sample characteristics (i.e., adults vs. children) and different acquisition protocols. This generalization may be attributed to the self-supervised nature of our method, which is optimized to learn effective representation of the intrinsic functional brain organization from input data without external guidance or prior assumptions. The robust generalization emphasizes that this approach is likely to be widely applicable to heterogeneous data in diverse applications.

Though the deep learning model is promising to learn highly reproducible and functionally homogenous FNs at individual level, there are still several potential limitations which should be noted. First, based on prior work, our method currently generates 17 networks. Future work will be devoted to evaluation of deep learning models for identifying FNs with different spatial resolutions. Second, as the human brain is a multi-scale system with hierarchical organization^41–49^, modeling personalized FNs at multiple scales with a hierarchical structure simultaneously could allow us to better account for the finding that hetero-modal association cortex in humans has multiple cognitive functions^50, 51^. Third, while we currently focus on identifying FNs for volumetric imaging data only, our method is directly appliable to cortical surface data by projecting the surface data to a 2D space^52^. Fourth, the functional network learning module adopts a U-Net architecture, which can be optimized using neural architecture search (NAS) techniques^53^.

Notwithstanding these limitations, we developed a self-supervised deep learning method to identify personalized functional networks accurately and efficiently. The evaluation of our method on multiple datasets has demonstrated that the deep learning model built using our method could identify functional networks in healthy adults, children, and patients with mental illness. Notably, personalized functional networks effectively captured individual differences in functional topography that could be used to predict development, cognitive performance, and schizophrenia status. Moving forward, this approach may be particularly useful for both large-scale functional imaging studies as well as smaller studies that target neuromodulatory interventions (e.g., TMS) using individual-specific functional neuroanatomy.

## ONLINE METHODS

### Imaging data

Four resting-state fMRI (rsfMRI) datasets from different cohorts were used to develop and validate the proposed method in the current study, including data from the Human Connectome Project (HCP), the Philadelphia Neurodevelopmental Cohort (PNC), the Pediatric Imaging, Neurocognition, and Genetics (PING) Data Repository, and one cohort of healthy controls and schizophrenia patients (SCZ).

#### HCP dataset

This dataset consisted of 1078 healthy individuals from the HCP cohort^29^. For each individual, two rsfMRI sessions (left-to-right direction phase encoding) were included. Details of the scanning protocols of the rsfMRI data has been published previously^54^. Frame-wise displacement (FD) was adopted to measure volume-to-volume changes in head position and participants with large head-movements (mean FD > 0.25 mm) were excluded. The ICA-FIX denoised rsfMRI in volumetric space were used, and the denoised rsfMRI data was spatially smoothed with a 6-mm full-width half-maximum (FWHM) kernel and downsampled at a spatial resolution of 3 × 3 × 3 *mm*^3^. The image data was downsampled to make its spatial resolution consistent with that of fMRI data from other 3 cohorts.

#### PNC

This dataset consisted of 969 young individuals (ages from 8 to 23 years, 535 females) from the PNC^34^. For each individual, a rsfMRI scan was acquired using 3T Siemens Tim Trio whole-body scanner with single-shot, interleaved multi-slice, gradient-echo, echo planar imaging (GE-EPI) sequence sensitive to BOLD contrast (TR = 3000 ms, TE = 32 ms; flip angle = 90°, FOV = 192 × 192 *mm*^2^, matrix =64 × 64, 46 slices, slice thickness/gap = 3/0 mm, effective voxel resolution = 3 × 3 × 3 *mm*^3^, 124 volumes/TRs). The rsfMRI data were preprocessed using an optimized procedure, including removal of the initial volumes, slice timing correction, 36-parameter confound regression, and band-pass filtering^55^. Participants with large head-movements (mean FD > 0.25 mm) were excluded.

#### PING dataset

This dataset consisted of 210 young individuals whose MRI scans were acquired using 3T Siemens scanners (ages from 3 to 20 years, 106 females) from the PING cohort^35^. For each individual, a rsfMRI scan with varied effective voxel resolutions of 4 × 4 × 4 *mm*^3^ or 3.75 × 3.75 × 3.5 *mm*^3^ was acquired using imaging sequence parameters optimized for equivalence in contrast properties and consistency in image-derived quantitative measures with integrated distortion correction. The rsfMRI data were preprocessed using the same procedure as the PNC dataset. Participants with large head-movements (mean FD > 0.25 mm) were excluded.

#### SCZ dataset

This cohort consisted of 195 individuals (101 healthy controls, and 94 SCZ patients) from previous studies^56, 57^. All the subjects underwent structural MRI and rsfMRI scanning with a 3.0-Tesla MR system (Discovery MR750, General Electric, Milwaukee, WI, USA). Tight but comfortable foam padding was used to minimize head motion, and earplugs were used to reduce scanner noise. Sagittal 3D T1-weighted images were acquired using a brain volume sequence (TR/TE/TI=8.2/3.2/450 ms; FA=12°; FOV=256×256 mm; matrix=256×256; slice thickness=1 mm, no gap; and 188 sagittal slices). A gradient-echo single-short echo planar imaging sequence was used to collect rsfMRI data (TR/TE=2000/45 ms; FOV=220×220 mm; matrix=64×64; FA=90°; slice thickness=4mm; gap=0.5mm; 32 interleaved transverse slices; and 180 volumes). All subjects were instructed to keep their eyes closed, relax, move as little as possible, think of nothing in particular, and not fall asleep during the rsfMRI scanning. All images were visually checked for artefacts, structural abnormalities, and pathologies by neuroradiologists. The rsfMRI scans were preprocessed using fMRIPrep^58^ (version: 20.2.1, https://github.com/nipreps/fmriprep/archive/20.2.1.tar.gz) and head motion artifacts were removed using ICA-AROMA^59^. Participants with large head-movements (mean FD > 0.25 mm) were excluded. The final dataset comprised data from 101 healthy controls (HC, age: 33.86 ± 10.77 years, 58 females) and 94 schizophrenia (SCZ, age: 33.23 ± 8.03 years, 41 females) subjects. There is no significant difference between the healthy controls and schizophrenia subjects regarding age (*p* = 0.647, two-sample T-test), gender (*p* = 0.0539, Chi-squared test), or head motion (*p* = 0.3069, two-sample T-test).

All the preprocessed rsfMRI data across datasets were in the MNI space and with spatial resolution of 3 × 3 × 3 *mm*^3^ so that the DL model worked with reception field of the same size (in convolutional operations) on all datasets.

### Functional brain decomposition based on matrix factorization

Given rsfMRI data *X^i^* ∈ *R*^*T*×*S*^ of individual *i*, consisting of *S* voxels and *T* time points, a matrix factorization based decomposition model tries to identify *K* FNs 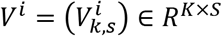 and their corresponding time courses 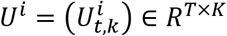, so that the original rsfMRI data can be approximated by two low-rank matrices *X^i^* ≈ *U^i^V^i^*. To identify FNs that do not contain anti-correlated functional units and are spatially compact and overlapped, non-negative constraint and spatial sparsity regularization are usually applied to *V^i^* ^10^, and the FNs can be obtained by optimizing

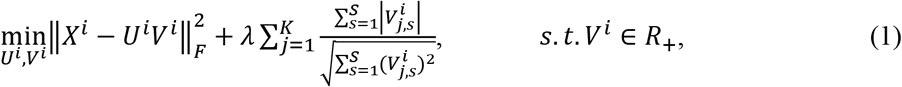

where the first term is a data fitting term, the second term is the Hoyer regularization for spatial sparsity^60^, and *λ* is the trade-off parameter.

An alternative optimization method is usually adopted to optimize Eq. (1)^61^. When *V^i^* is fixed, *U^i^* can be calculated analytically as *U^i^* = *X^i^*(*V^i^*)^*T*^(*V^i^*(*V^i^*)^*T*^)^−1^. Substituting this expression for *U^i^* in Eq. (1), we have

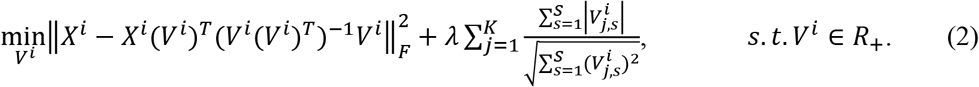

Instead of optimizing *V^i^* by the multiplicative weight update method^61^, we propose to train a deep convolutional neural network of CNNs to estimate *V^i^* directly from the input rsfMRI data as shown in Fig. 1a.

### Self-supervised deep learning of personalized FNs

Personalized FNs are computed from rsfMRI using CNNs with an Encoder-Decoder architecture under a self-supervised deep learning framework by optimizing data fitting and sparsity regularization terms that are commonly used in brain decomposition models^10, 36^. A trained model can be applied to rsfMRI data of a new individual to identify personalized FNs in one forward-pass computation. As illustrated in Figure 1, we adopt an Encoder-Decoder U-Net^62^ to identify personalized FNs from rsfMRI data by optimizing data fitting and spatial sparsity regularization terms^10, 36^ so that the network is trained and optimized based on the input rsfMRI data without any external supervision. The Encoder-Decoder CNNs facilitate the inter-individual comparability of FNs implicitly by an effective representation of intrinsic FNs in a low-dimensional latent space characterized by the Encoder-Decoder’s bottleneck layer.

Given a group of *n* individuals, each having 4D rsfMRI data *X^i^* ∈ *R*^*W*×*H*×*D*×*T*^, *i* = 1,…, *n*, where *W*, *H*, and *D* are width, height, and depth of each 3D volume respectively and *T* is the number of time points, we train a deep network of 3D CNNs *M_θ_c__*(*I*) = *V_I_* with convolutional parameters *θ_c_*, which takes the 4D fMRI data *I* as input and identifies its corresponding FNs *V_I_* as output. We use rsfMRI data from *n* individuals as training data to minimize the loss function in Eq. (2) for learning the convolutional parameters *θ_c_* (when calculate the loss function, *I^i^* is flatten across the spatial dimension, transposed and reshaped as a matrix with size *T* × *S*, where *S* = *W* × *H* × *D*. The same transformation is applied to *V_I_* to get the reshaped matrix with size *K* × *S*). As FNs are optimized as formulated by Eq. (2), which does not rely on any external supervision, the deep learning model is optimized based on input data alone, referred to as self-supervised. Once *θ_c_* is optimized, the deep learning model can be used to predict personalized FNs for different individuals.

The deep network includes a representation learning module and a functional network learning (U-Net^62^ like) module for identifying FNs, as illustrated in Fig. 1a. The representation learning module (with details in next paragraph) extracts time-invariant feature maps from the input rsfMRI using 3D CNNs, and the feature maps are used as input to the functional network learning module. As shown in Fig. 1c, the functional network learning module consists of one convolutional layer with 16 filters, three convolutional layers with 32 filters and a stride of 2, three deconvolutional layers with 32, 32, and 16 filters and a stride of 2, and finally 2 convolutional layers with 16 filters. Leaky ReLU activation and batch normalization^63^ are used for all the convolutional and deconvolutional layers. One output convolutional layer is used to predict personalized FNs and the number of output channels is *K*. Rectified linear activation function (ReLU) activation is used for the output layer so that the output is non-negative. Each output channel is linearly scaled so that its maximum equals to 1 before calculating the loss function. The kernel size in all layers is set to 3×3×3.

Inspired by a prior study^64^ which demonstrated that replication of FNs can be obtained through averaging of selected fMRI time frames, we propose to learn features from each individual time point of rsfMRI scans and fuse the features of all time points for learning personalized FNs by the following functional network learning module. As shown in Fig. 1b, the representation learning module consists of two parts referred as temporally separated convolution and temporal feature fusion. The temporally separated convolution part is a 3D convolutional layer with *C* filters and a stride of 1, which is applied to each time point (a 3D volume with size of *W* × *H* × *D*) of the rsfMRI scan and output *C* feature maps (with size of *W* × *H* × *D* × *C*) for each time point. The temporal feature fusion part is an element-wise average pooling layer which outputs *C* average feature maps of all the time points. In the present study, *C* is set to 16 and the kernel size of the convolutional layer is set to 3×3×3, and Leaky ReLU activation and batch normalization are also adopted. While the convolution learning part is expected to capture the co-activation patterns of different brain regions by different feature channels at each time point, the feature fusion part summarizes the patterns obtained from all the time points. This module is plugged into the model just before the functional network learning module, and the whole model can be optimized in an end-to-end fashion.

In the present study, the number of FNs is set to 17 in order to facilitate a direct comparison with the well-established FNs^30^. Our model is implemented using Tensorflow. Adam optimizer is adopted to optimize the network, the learning rate is set to 1 × 10^−4^, the batch size is set to 1, and the number of iterations is set to 30000 during training. One NVIDIA TITAN Xp GPU is used for training and testing. We set *λ* = 10 empirically in our experiments.

### Functional homogeneity

For each FN, a homogeneity measure is calculated as the weighted sum of the correlation coefficients between the time courses of all the voxels within the FN and its centroid time course, which is a weighted mean time course within the FN and the FN’s voxelwise loadings are used weights. The median value of FN-wise homogeneity measures is used to gauge functional homogeneity of FNs for each subject.

### Sanity testing for personalized FNs computed by the proposed DL model

Sanity testing was performed for quality assurance of the personalized FNs in terms of both functional homogeneity and spatial correspondence. First, the personalized FNs was compared with their corresponding group average FNs in terms of within-network functional homogeneity. Specifically, given a subject with preprocessed fMRI data, functional homogeneity measures are computed based on its personalized FNs and the group average FNs. If a subject’s personalized FNs have higher functional homogeneity than the group average FNs, then the sanity testing is passed in terms of the functional heterogeneity. Second, the personalized FNs’ spatial correspondence across individuals is tested. Instead of comparing FNs across individuals directly, each individual subject’s FNs are compared with group average FNs based on their spatial correlation coefficients. Specifically, one personalized FN is deemed to maintain correspondence with its corresponding group level FN if 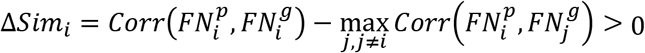, where 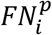 denotes the *i*-th personalized FN, 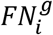 denotes its corresponding group average FN, 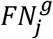 denotes other group average FNs, and *Corr*(·,·) is spatial correlation coefficient between two FNs. If a subject’s all personalized FNs maintain the correspondence with their group average FNs, then the sanity testing is passed in terms of the spatial correspondence.

### Ridge regression

To investigate if personalized FNs identified by the DL model were related to behavior and development, we used ridge regression to predict 13 cognitive measures (using data from HCP) and brain age (using data from PNC and PING) at individual level using the network loadings from all 17 FNs as features in a multivariate regression setting. Specifically, we used ridge regression as implemented in scikit-learn^65^. The prediction was performed under a 2-fold cross-validation setting, and the regularization parameter *λ* was identified in the range [2^−10^, 2^−9^,…, 2^4^, 2^5^] using nested 2-fold cross-validation within the training fold. The optimal *λ* was used to train the prediction model on the training fold, which was then applied to the testing fold for evaluation.

### SVM classification

To investigate if personalized FNs identified by the DL model were related to neuropsychiatric illness, we used support vector machine (SVM) classifier to distinguish patients with schizophrenia from healthy individuals using the network loadings from all 17 FNs as features. Specifically, we used SVM with a linear kernel as implemented in scikit-learn^65^. The classification was performed under a 2-fold cross-validation setting, and the regularization parameter *C* was identified in the range [2^−10^, 2^−9^,…, 2^4^, 2^5^] using nested 2-fold cross-validation within the training fold. The optimal λ was used to train the prediction model on the training fold, which was then applied to the testing fold for evaluation.

## SUPPLEMENTAL INFORMATION

**Fig. S1.**
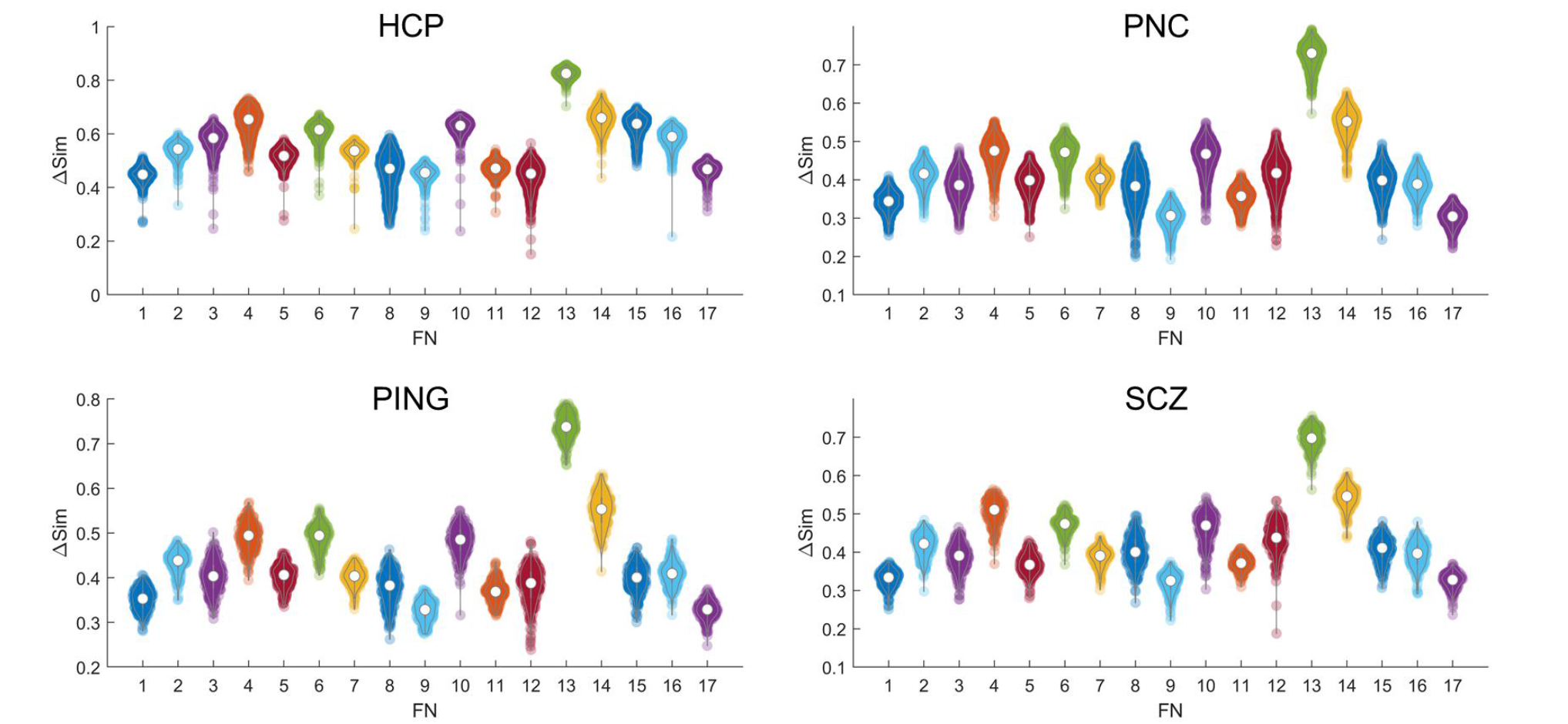
Spatial correspondence of personalized FNs are well preserved for testing individuals across different cohorts (HCP, PNC, PING, and SCZ). The spatial similarity of each personalized FN and its corresponding group FN is clearly higher than its similarity with all other group FNs (all Δ*Sim* > 0.1). In each plot, one datapoint denotes the Δ*Sim* of one FN from one testing individual.

**Fig. S2.**
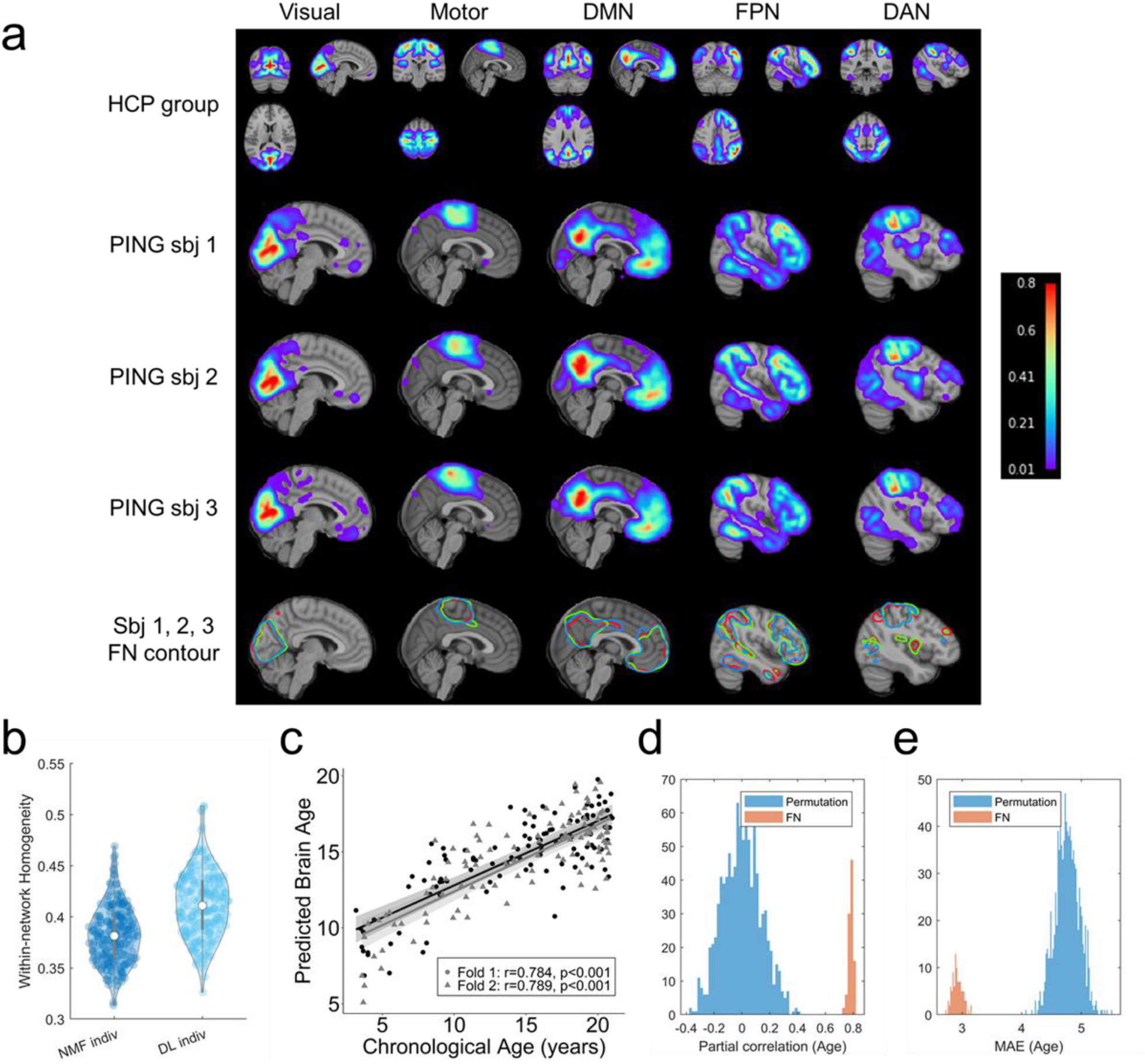
The DL model trained on the HCP cohort generalizes well to the PING cohort. **(a)** Personalized FNs of three randomly selected PING subjects, identified by the DL model trained on HCP cohort, in sagittal view (the second, third, and fourth rows), isolines of the FNs at a value of 0.15 in different colors (the fifth row), and their corresponding average FNs of all testing HCP subjects (the first row). **(b)** Personalized FNs computed using the DL model had significantly higher functional within-network homogeneity than those computed using the spatially-regularized NMF (*p* < 10^−5^, Wilcoxon signed rank test). **(c)** Age prediction performance (partial correlation coefficients) of the personalized FNs, obtained with one run of the 2-fold cross-validation. Data points in different colors represent the prediction results of different folds in the 2-fold cross-validation. **(d, e)** Prediction accuracy measured with partial correlation coefficients and MAE values of 100 runs of the 2-fold cross-validation and the null distribution of prediction accuracy from permutation tests (1000 runs of 2-fold cross-validation on permuted data).

